# Repeated and widespread introduction of a mosquito-borne virus, but wildlife impact remains localised in urban area

**DOI:** 10.64898/2026.02.03.703450

**Authors:** Hugh J. Hanmer, Robert C. Bruce, Ava J. Jenkins, Mariana Santos, Katharina Seilern-Macpherson, Sarah Haddow, Emily Jones, Shinto K John, Andra-Maria Ionescu, Bathsheba L. Gardner, Anthony J. Abbott, Nicholas Johnson, Alexander G.C. Vaux, Simon Spiro, Andrew A. Cunningham, Jolyon M. Medlock, Becki Lawson, Robert A. Robinson, Arran J. Folly

**Affiliations:** British Trust for Ornithology, The Nunnery, Thetford, Norfolk, IP24 2PU, UK; Animal and Plant Health Agency, Woodham Lane, Addlestone, Surrey, KT15 3NB, UK; Institute of Zoology, Zoological Society of London, Regent’s Park, London, NW1 4RY, UK; Medical Entomology and Zoonoses Ecology Group, UK Health Security Agency, Salisbury, SP4 0JG, UK; Wildlife Health Services, Zoological Society of London, London, UK

**Keywords:** Usutu virus, flavivirus, Eurasian Blackbird, climate, emerging infectious diseases, vector-borne disease, zoonosis

## Abstract

Mosquito-borne viruses are emerging in new regions with increasing regularity. Shifting climatic envelopes, especially in temperate regions, and altered enzootic cycles are changing likelihoods of persistence, meaning new approaches are required to predict and monitor establishment. Usutu virus (USUV) is a mosquito-borne viral zoonosis, which was first recorded in the United Kingdom in wild birds and mosquitoes in 2020. By developing a multidisciplinary approach, incorporating enhanced passive and active surveillance of wild birds, we discovered previously unrecorded introductions and identified USUV persistence and geographic expansion. By combining molecular and serological testing with citizen science derived datasets, we show an associated marked decline in a common, widespread, and highly susceptible host species (Eurasian Blackbird, *Turdus merula*). However, the clearest continuous signal of host population decline remains localised to a large urban area, despite the much wider virus distribution. This may be a consequence of an urban heat island effect and associated increase in length of the mosquito active season. Our results indicate that the establishment and impact of future mosquito-borne pathogens in temperate zones, anticipated under climate change, may first be most apparent in urban areas. Consequently, enhanced surveillance efforts in urban areas could provide an efficient mechanism for detecting novel pathogen emergence. The epidemiology of emerging vector-borne diseases, especially in temperate areas, depends on the interplay between climate, land-use and available hosts; therefore, considering each in isolation will not be sufficient to reliably inform surveillance or response planning.

## Introduction

Emerging mosquito-borne viruses are causative agents of disease outbreaks impacting both animals and humans [1-3]. However, determining when they become established, and quantifying impact on susceptible hosts, can be difficult [4]. This is often the case in temperate regions, as mosquito-borne viruses require specific climatic envelopes and vector-host enzootic cycles, which may not exist sufficiently to facilitate persistence [5], or may result in undetected, low-level, yet sustained, transmission [6]. Consequently, without adequate surveillance, novel introductions and geographic spread may be underestimated or overlooked prior to notable disease outbreaks in susceptible hosts [6].

Usutu virus (USUV) is a mosquito-borne viral zoonosis [7, 8], which was first detected in southern Africa in 1959, before emerging in Europe during the 1990s [7]. The virus exists in an enzootic cycle between mosquitoes (typically from the *Culex* genus) and wild birds. Some species (e.g. *Culex modestus*), can act as bridge vectors transmitting the virus to non-target animals, including humans, where infection may result in neurological disease [9]. However, passerines (especially Eurasian Blackbird [*Turdus merula*]) and owls (Strigiformes) are highly susceptible hosts [10, 11] [12]. The geographic range of USUV is presumed to expand through the movement of vectors and viraemic hosts. Expansion may be over relatively short distances (< 10km), facilitated by movement of mosquitoes [13] or territorial hosts, but can occur over much larger distances, through the migration of viraemic birds [14].

Usutu virus is now considered endemic in south-eastern England, since its first detection in wild birds in the United Kingdom in 2020 [15, 16]. Seroconversion subsequent to the index USUV case in London has been recorded in a range of exotic bird species housed at ZSL London Zoo [16] and annual detection of USUV RNA, primarily from Blackbirds [17], indicated sustained transmission at the index site [16], which was associated with a local decline in Blackbird numbers [18]. To improve our understanding of how orthoflaviviruses emerge, spread and impact their hosts, we adopted a multidisciplinary approach to investigate USUV infection in birds in the UK, which comprised both enhanced passive and active surveillance approaches. In addition, we interrogated citizen science derived datasets to elucidate any impact of USUV infection on susceptible hosts. Combined, these approaches provided evidence for multiple introductions, geographic spread with continued USUV persistence, while simultaneously implicating a large urban area in facilitating transmission of USUV and associated mortality in susceptible wild bird hosts.

## Methodology

### Passive surveillance of birds

We enhanced our passive surveillance, by combining existing national scanning surveillance schemes [18] with a risk-based approach using a sentinel species. Samples from a total of 1,391 wild and captive birds of 108 species were obtained for passive *Orthoflavivirus* surveillance (figure 1). Brain and kidney samples were collected at post-mortem examination (PME) from wild birds submitted from across Great Britain and from captive birds that died at ZSL London Zoo and Whipsnade Zoo during the peak UK mosquito active season, April-November inclusive, in 2023 and 2024 (supplementary table 1). Brain and kidney samples were pooled for all downstream molecular analysis. Available PCR-positive tissues were subjected to immunohistochemical analysis to detect flavivirus envelope antigen, as previously described [18].

**Figure 1.**
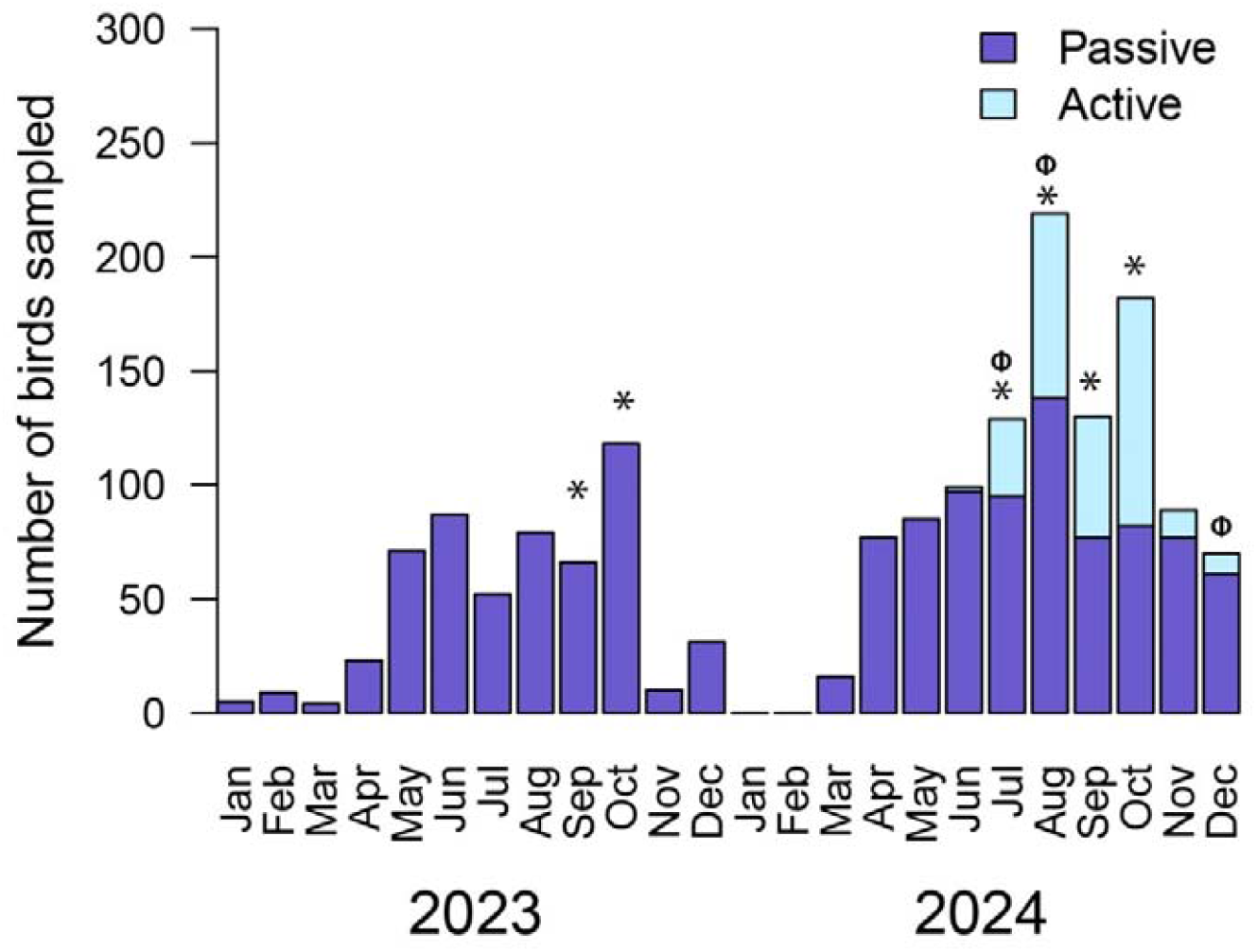
Active (2024) and enhanced passive surveillance in birds for 2023 and 2024 across Great Britain. The ^*^ indicate months with USUV RNA detection, and □ indicate months with detection of *Orthoflavivirus* seroconversion.

Blackbird was selected as a sentinel species for USUV detection as it is a common, USUV-susceptible species that is frequently admitted to wildlife rescue centres. Using a risk-based approach, focused on the geographic region predicted to have the greatest likelihood of USUV circulation, five wildlife rescue centres in southern England were recruited. Blackbirds that died, or were euthanised for welfare reasons while in care, were stored at - 20×C pending collection and PME. A total of nine Blackbirds that died between July and October 2023, and 52 Blackbirds that died between July and September 2024 inclusive, were examined post mortem using previously-published methods [19]. Along with brain and kidney samples, primary flight and body contour feathers and any attached ticks, were collected from Blackbirds for PCR testing. Samples were stored at -80×C.

### Active surveillance of passerines

Live wild passerines (n = 292) were sampled under Home Office Licence PP9908514 by personal licensees between July and December 2024, inclusive (figure 1). To elucidate viral incursion and persistence dynamics, samples were obtained from a typically resident UK species (House Sparrow (*Passer domesticus*) [n = 23]), partial migrant species (Blackbird [n = 51], Song Thrush (*T. philomelos*) [n = 17]) and migratory species (Blackcap (*Sylvia atricapilla*) [n = 85], Chiffchaff (*Phylloscopus collybita*) [n = 75], Common Whitethroat (*Curruca communis*) [n = 12], Lesser Whitethroat (*Curruca curruca*) [n = 2], and Willow Warbler (*Phylloscopus trochilus*) [n = 27]). The latter group typically spend their non-breeding periods in Africa and the Mediterranean region where West Nile virus (WNV) and USUV circulate. A risk-based approach focussing on south-eastern England was adopted to maximise the likelihood of USUV detection. Each individual was sampled during bird ringing activities and checked to make sure they were healthy before being sampled. Sampling comprised manual contour feather removal (n ≤ 5 feathers stored dry in 2ml Eppendorf tube), separate 4mm oral and cloacal swabs (ClassiqSwabs, Italy), with molecular grade water moistened tips, stored in RNAlater (Invitrogen, USA), and a blood sample. Blood sampling technique differed depending on bird body weight to ensure that no more than 10% of circulating blood volume was removed for any bird, which is the accepted safe limit (www.nc3rs.org.uk). For the majority of birds ≥ 18.2g, an active blood draw from the right jugular or brachial vein was performed using a sterile 0.5ml insulin syringe (Becton Dickinson, USA) to remove ≤ 100µl of circulating blood and stored in a serum gel Minicollect ® tube (Greiner bio one, USA). For birds ≤18.2g, a brachial vein prick (≤ 20µl) using a 30-, 27- or 25-gauge sterile hypodermic needle was done, followed by taking a blood spot on protein preservation card (903™ Protein saver card, Whatman, USA). Immediately after blood collection, direct pressure was applied on the puncture site with cotton wool until haemostasis was achieved. Following sampling, a final health assessment was performed by a ringing permit holder or veterinarian before release. No adverse effects of sampling were recorded.

### Molecular surveillance and phylogenetic analysis

Total RNA was extracted using RNeasy Mini-Prep (Qiagen, UK), following the manufacturer’s protocol. Extracted RNA was eluted in 50µl of RNase free water. All samples were subjected to USUV-specific RT-PCR assay using published protocols [20]. Amplicons of the anticipated size were submitted for confirmatory Sanger sequencing. For those returning USUV sequence, the original RNA sample was subjected to next generation sequencing (NGS). Sequencing libraries were prepared using the Nextera XT kit (Illumina, Cambridge, UK) and analysed on a NextSeq sequencer (Illumina, Cambridge, UK) with 2 x 150bp paired-end reads. Consensus sequences were generated by using a combination of the Burrows–Wheeler Aligner v0.7.13 and SAMtools v1.9 with a representative USUV genome (GenBank accession number: MW001216) as a scaffold. Contiguous sequences obtained through NGS were aligned with representative USUV isolates downloaded from GenBank (supplementary table 2) using MAFFT v7.471. The resulting alignment was imported into BEAST (v1.10.4) and a Bayesian phylogenetic tree was produced using the GTR+I nucleotide substitution model and 10,000,000 Markov chain Monte Carlo generations. Log files were analysed in Tracer v1.7.1 to check the effective sample size and a 10% burn-in was included (TreeAnnotator v.1.10.4) before visualisation and annotation in FigTree v1.4.4.

### Serological testing

Whole blood samples from wild birds were centrifuged to separate serum, whereas blood samples on protein preservation cards were resuspended in 150µl of 1% phosphate buffered saline (Sigma Aldrich, UK). For each sample, 25µl was screened for anti-*Orthoflavivirus* activity using a commercially available enzyme-linked immunosorbent assay (ELISA) (IDvet, Grabels, France). For ELISA-positive samples, if there was sufficient serum remaining, a plaque reduction neutralisation test (PRNT) was undertaken on 12-well plates seeded with ≥80% confluent Vero cells. Samples were serially diluted (1:10,1:20 etc) in E-MEM (Thermo-Fisher, UK) and combined with USUV (lineage 3.2, GenBank accession number: MW001216) to confirm antibody specificity. Two millilitres of carboxy-methylcellulose was used to cover each well, and plates were incubated for 4 days at 37×C and 5% CO_2_, before fixation and staining using crystal violet (Sigma-Aldrich, UK). Serum that neutralised USUV at or above a plaque reduction threshold of 90% was considered to contain USUV-specific neutralising antibodies.

### Elucidating the impact of Usutu virus infection on Blackbird population

The British Trust for Ornithology (BTO) Garden BirdWatch (GBW) is a garden wildlife citizen science survey, with approximately 6,000 volunteers systematically submitting data for the presence and abundance of all observed bird species in their gardens each week throughout the calendar year across the UK [21].

To investigate the change in frequency with which Blackbirds are reported in gardens over time, weekly reporting rate of Blackbirds was summarised (i.e. proportion of gardens recording the species) for each GBW garden during the breeding period for each year 2017-2024 inclusive. The breeding period was defined as calendar weeks 13 to 19 inclusive (approximately 26^th^ March – 13^th^ May). This encompasses the timing of the majority of first nesting attempts [22] and represents a period when the proportion of gardens from which Blackbirds are reported is consistent across the UK (∼95% in 2017; supplementary figure 1). To ensure sufficient coverage, we considered only GBW gardens (range: 4965 - 7088 gardens/year) that were monitored for at least two years during the eight-year study period, and years in which data were submitted in at least 5 of the 7 weeks of the breeding season. The proportion of active weeks in the breeding period for which Blackbirds were absent for each season and GBW garden was calculated and then summarised as the mean proportion of weeks absent for each UK Ordnance Survey 10 km grid square that contained a minimum of two gardens in any given year.

## Results

### Passive surveillance of wild birds for USUV

Samples from fifteen birds tested positive for USUV RNA (supplementary table 3) comprising ten Blackbirds (six of which were from rescue centres), one Feral Pigeon (*Columba livia*), and four collection birds from ZSL London Zoo (two Great Grey Owls (*Strix nebulosa*) and two Grosbeak Starlings (*Scissirostrum dubium*)). Seven of the eight PCR positive Blackbirds for which fixed tissues were available had microscopic abnormalities and flavivirus envelope (E) antigen in assessed tissues (supplementary table 4), consistent with systemic USUV infection as the probable cause of death. Trauma was the cause of death of the remaining Blackbird with absent flavivirus labelling.

On phylogenetic analysis, five Blackbirds (from Buckinghamshire, Cambridgeshire, Greater London, Hertfordshire and Oxfordshire), and the four zoological collection birds from Greater London, were identified as being infected with USUV lineage Africa 3.2, most closely related to previous UK USUV detections (GenBank accession numbers: MW001216 & OM202464). Three other Blackbirds from the same counties were characterised as USUV lineage Africa 3.2, however, we obtained insufficient genome coverage for inclusion in phylogenetic analysis. Two Blackbirds, one from East Sussex (50.94° N, 0.27° E) (USUV lineage Africa 3.1 GenBank accession number PX776633) and one from Bedfordshire (50.06° N, 0.52° W) (USUV lineage Africa 3.2 GenBank accession number PX776632), were infected with lineages of USUV that were previously undetected in the UK (figure 2). The PCR positive Feral Pigeon was infected with USUV Africa 3.2.

**Figure 2.**
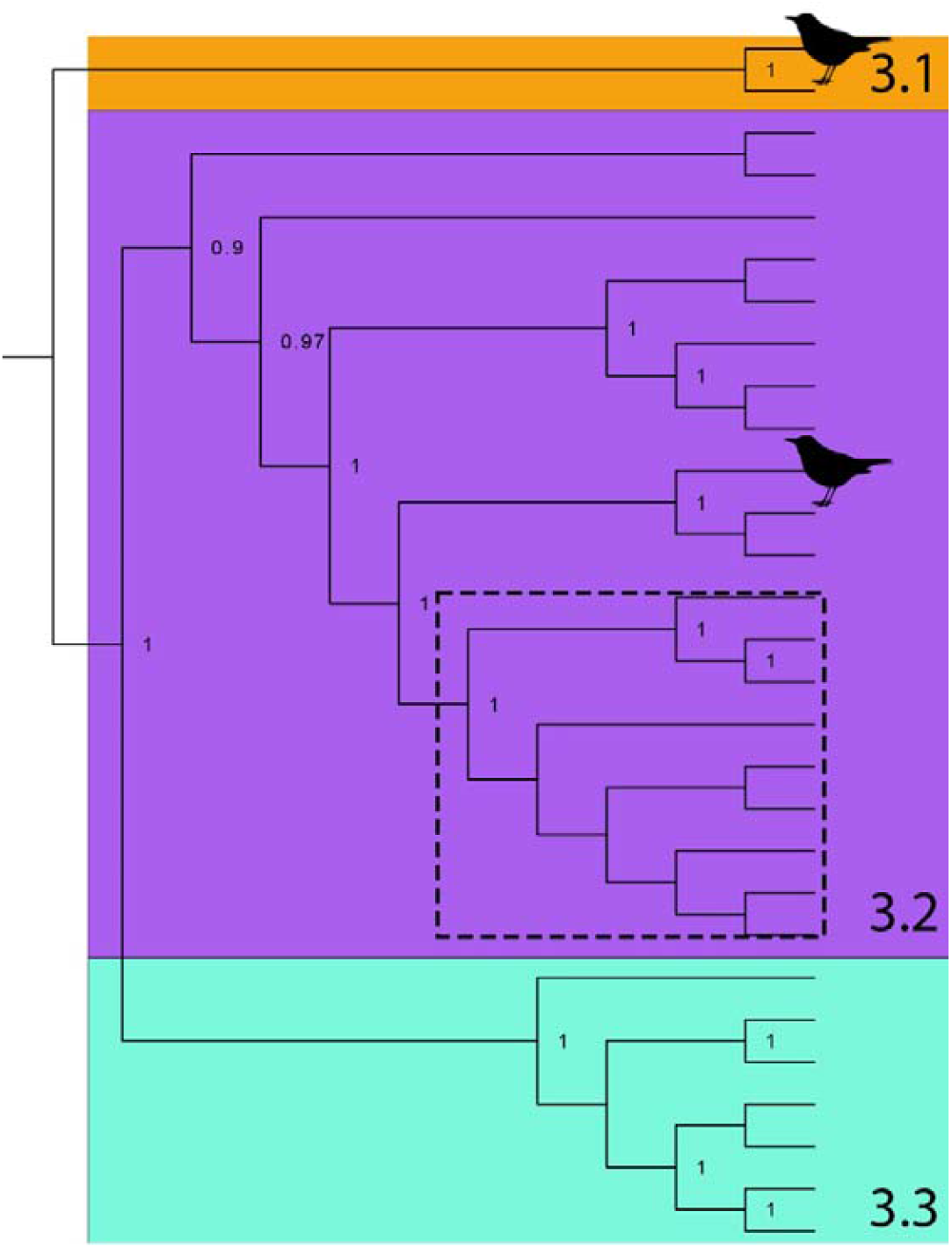
Bayesian phylogeny of USUV Africa lineages, based on 9997 base pairs showing previously unidentified USUV lineages from Blackbirds (Blackbird icon) in south-eastern England. Hashed box indicates all previous UK USUV detections (GenBank accession numbers PX776632 & PX776633). Posterior probability of nodes indicated where greater than 0.75.

### Active surveillance of wild passerines for USUV

USUV lineage Africa 3.2 RNA was detected in eight birds (2.7%) from across our sampling area (supplementary table 3). These birds comprised juvenile (hatched that calendar year) Blackcap (n=3), Chiffchaff (n=4) and Common Whitethroat (n=1), which are all migratory species in the UK. A 150bp PCR amplicon, from a Chiffchaff oral swab sample collected in Kent (51.09° N 0.84° E), matched USUV lineage Europe 3 (through GenBank BLAST), although the quality of the sequence was insufficient to deposit on GenBank.

Eight birds, from across the sampling range, had evidence of *Orthoflavivirus* seroconversion (2.89% of 276 blood-sampled birds). Of these, four were Blackbirds (three adults (i.e. in at least their second calendar year, based on plumage) and one of undetermined age). No bird with confirmed USUV RNA (n = 8) had evidence of seroconversion. All detections through active surveillance were from areas where there was no prior knowledge of USUV circulation in the UK (figure 3).

**Figure 3.**
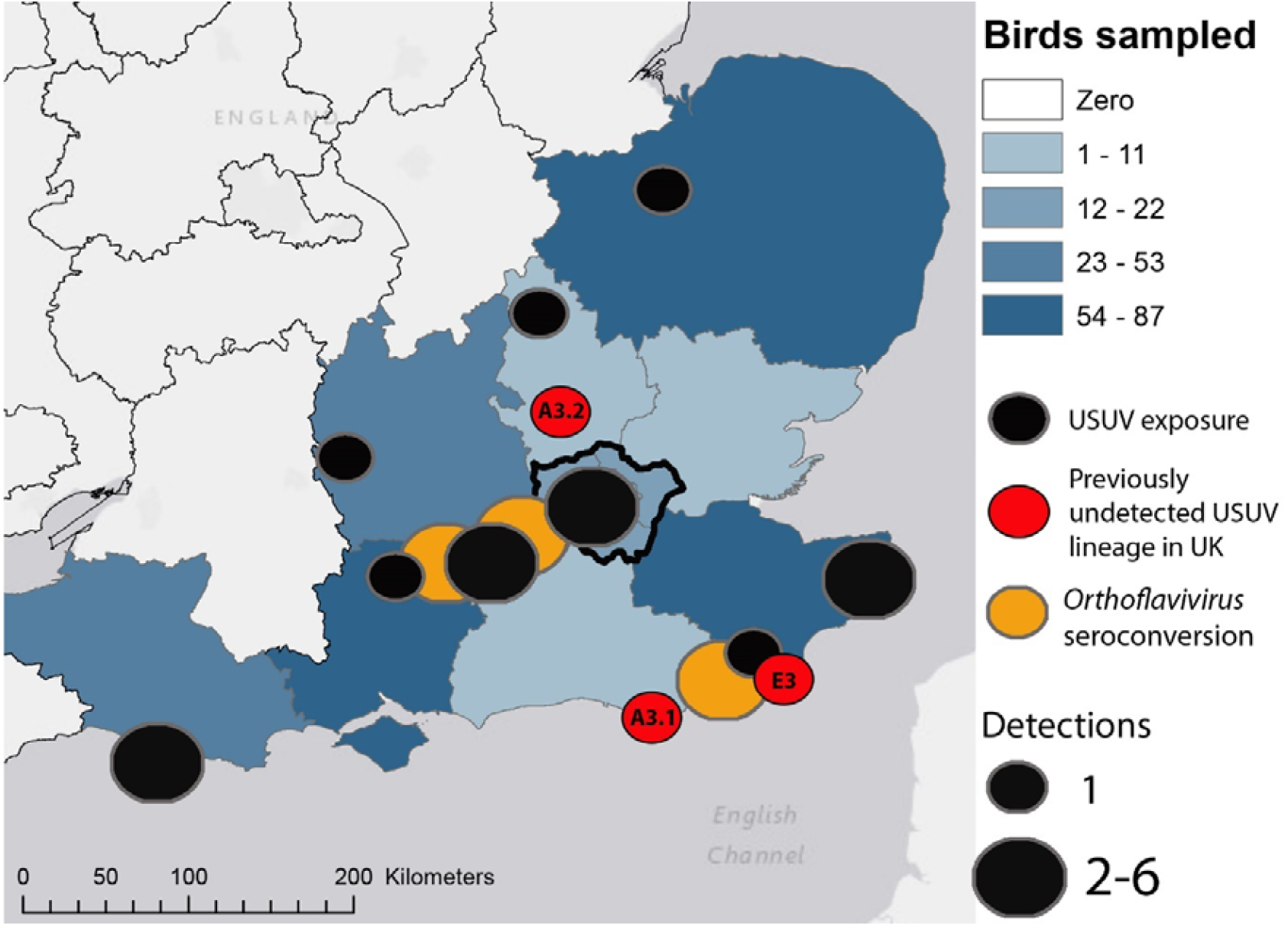
Active (2024) and enhanced passive (2023-2024, inclusive) *Orthoflavivirus* surveillance effort in birds from south-eastern England, shown by county. Black and red icons indicate sites with evidence of USUV RNA, with red icons indicating three USUV lineages previously unidentified in the UK (Africa 3.1 [A3.1], Africa 3.2 [A3.2] and Europe 3 [E3]), and orange icons indicate *Orthoflavirus* seroconversion. Dark border defines Greater London region.

### Elucidating the impact of Usutu virus infection on Blackbird population

Blackbirds were recorded in most gardens and weeks across the UK (mean >99% of gardens and 95% of weeks) during the 2017-2020 breeding seasons (supplementary figure 2), although marginally less frequently in Greater London (mean ∼98% of gardens and 83% of weeks, figure 4). There was a noticeable decline in the occurrence of Blackbird in Greater London during subsequent breeding seasons (2021: 80% of gardens and 56% of weeks; 2022: 78% and 48%; 2023: 64% and 35%; 2024: 68% and 38%). While the affected area increased, particularly in 2023 and 2024, it remained much smaller than the area from which positive USUV samples were obtained (figure 3). Only Blackbird appears to have been affected, as broadly ecologically similar [18] and USUV-susceptible species [23] showed no comparable declines in GBW recording rate in Greater London (supplementary figure 3).

**Figure 4.**
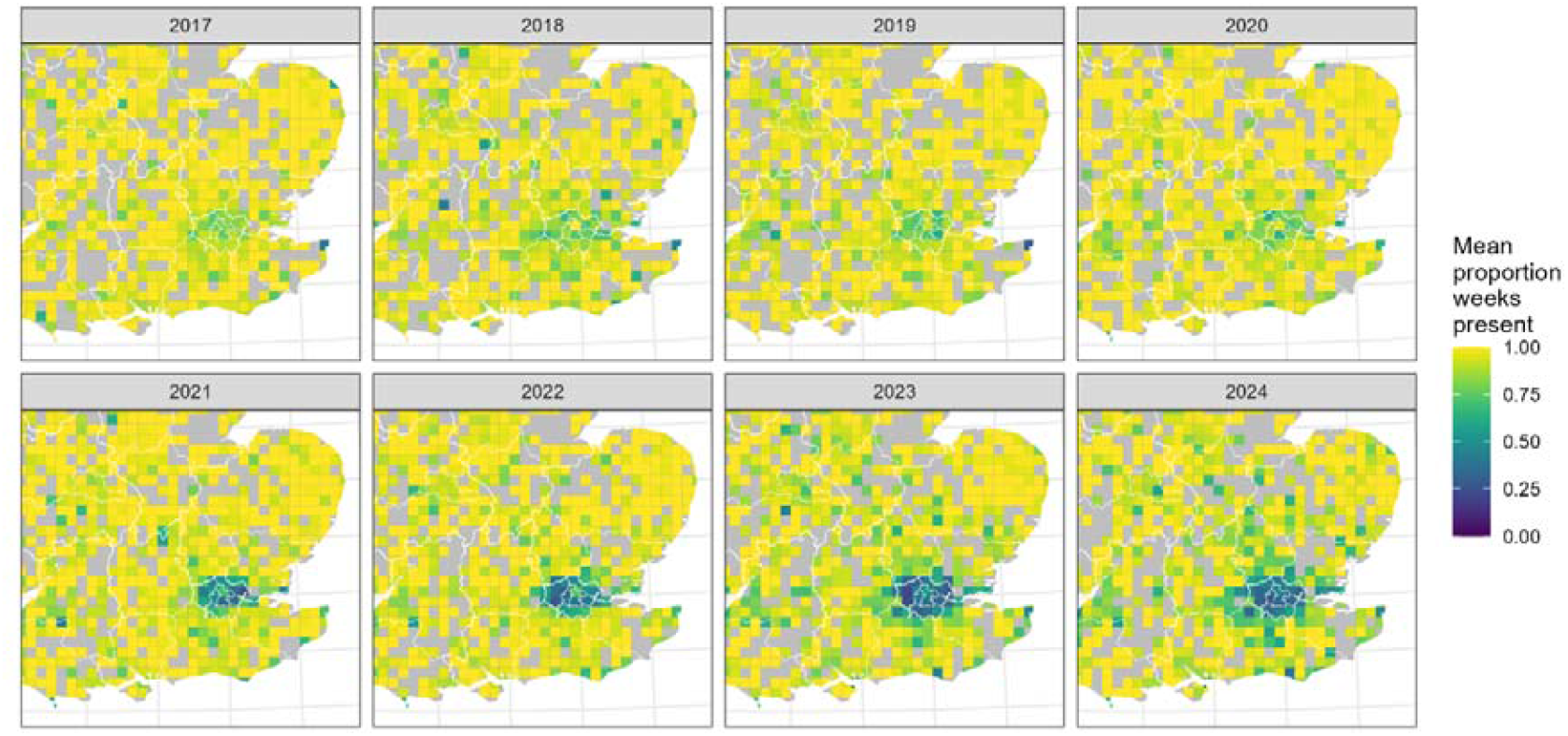
Maps showing change in the mean proportion of weeks when Blackbirds were recorded in British Trust for Ornithology’s Garden BirdWatch survey participant gardens per 10km grid square in south-eastern England on an annual basis, over the period 2017-2024. Grey indicates fewer than two gardens fulfilling the criteria, defined in methods, were active in that square in that year.

## Discussion

Our results indicate USUV is markedly more widespread in south-eastern England than previously recognised [17], and that multiple introductions of USUV have occurred. USUV infection appears to be the principal driver of Blackbird population decline, as has been observed in mainland European countries [10]. However, our results show that impacts on Blackbird populations are most notable in the Greater London region. We speculate that this could be related to the different environmental conditions encountered in large urban areas, which may support viral transmission in temperate zones [24-26].

USUV has been recorded annually in wild and/or captive birds in the UK since its first detection in 2020 [15, 16]. Prior to 2023, all identified isolates were most closely related to each other and to the original USUV Africa 3.2 UK detection, indicative of establishment and overwintering from a single introduction event [16, 17]. However, we obtained evidence that multiple introductions of USUV into the UK have occurred. These detections increase the known geographic range of USUV in England and highlight the continued risk of mosquito-borne *Orthoflavivirus* introductions to temperate countries [5]. During our surveillance we only detected single cases of infection with Africa 3.1 and Europe 3, indicating that these lineages may not be widely circulating. We also recovered USUV RNA from *I. frontalis* ticks attached to an infected Blackbird, providing evidence that they may be involved in transmission cycles [27]. More broadly, these results indicate that the UK may be susceptible to additional incursions from emerging mosquito-borne viruses, likely driven by the movement of viraemic avian hosts [28, 29] and/or mosquito vectors [30-32].

Active surveillance highlighted that a range of birds have been exposed to USUV. Specifically, USUV RNA was detected in samples collected from migratory species from across our sampling range. Interestingly, these birds were all identified as hatched within the calendar year in which they were sampled, therefore it is likely that infection occurred locally in the UK. In addition, USUV RNA was detected in samples from migratory birds that were collected during October 2024, from locations on the south coast of England, indicating they were commencing their southbound migration to non-breeding grounds and therefore also likely contracted USUV infection in the UK [33]. We were unable to isolate virus from the Chiffchaff oral swab and attempts at Sanger sequencing using different PCR products [34] were unsuccessful, however the initial result indicating USUV lineage Europe 3, suggests a previously undetected introduction of this lineage to the UK. Combined, our results highlight a dynamic network of *Orthoflavivirus* transmission, possibly with infected birds moving into and out of the UK [35].

We detected *Orthoflavivirus*-specific antibodies in migratory, partially migratory and resident bird species, with no evidence of USUV RNA in concurrent swab or feather samples. As most of these individuals were juvenile and, given the time of year, presumably hatched in the UK, it is probable they were exposed to USUV locally in the UK and acquired some immunity to infection through USUV-specific seroconversion [36, 37]. However, if birds were sampled shortly after fledging, we note that *Orthoflavivirus* antibodies may have been maternally derived [38].

While none of the active surveillance Blackbird samples indicated the presence of USUV RNA, four Blackbirds were positive on both ELISA and PRNT for USUV-specific antibodies. Four other birds tested positive on the ELISA, but the remaining serum was insufficient to conduct PRNT and thus confirm antibody specificity. Cross-reactivity among orthoflaviviruse*s* could provide false USUV-positive results using the ELISA alone. Cross reactive flaviviruses detected in the UK include USUV, WNV, louping ill virus (LIV) and tick-borne encephalitis virus (TBEV). While LIV is well established in the UK it has not been reported in passerines, similarly TBEV specific antibodies have, so far, only been detected in Blackbirds in mainland Europe [39]. All the Blackbirds for which USUV specificity was confirmed on PRNT were adults, and no USUV RNA or signs of disease were observed in these, indicating that some Blackbirds can mount a protective immune response following USUV exposure. None of the juvenile Blackbirds we sampled had evidence of seroconversion. The majority of the seroconverted Blackbirds were sampled during December 2024, which is outside the typical ornithophagic mosquito-active season in the UK (April-November, inclusive) [40]. This indicates that adult Blackbirds maintain seropositive status, which may facilitate some tolerance or resistance to USUV infection, although this effect may not be as apparent in migratory Blackbirds [41]. However, some resistance may be developing in adult Blackbirds which, in time, could facilitate recovery of the Greater London population, as has been observed in mainland Europe [11, 42]. Thus, by combining active and enhanced passive surveillance with citizen science derived population monitoring, we highlight that while there have been multiple incursions of USUV across south-eastern England, disease impacts at a population scale in Blackbirds appear so far to be localised, and that urban areas may facilitate transmission cycles of mosquito-borne viruses.

While Greater London was the index site for the UK USUV outbreak, we recognise it is unlikely to be the exact location of viral emergence [43]. Our results indicate that since the first detection of USUV in the UK, Blackbird populations in Greater London have declined [44] but we do not see the same signal in other regions where we have detected the virus. Given that urban areas are often warmer, with more stable climates [24] it is likely that in temperate regions, urban areas may facilitate arboviral establishment [3, 45, 46]. Indeed, warmer conditions can reduce viral extrinsic incubation periods [47] while simultaneously increasing the length of the mosquito active season [40], resulting in a greater proportion of the year when mosquito-borne viruses could be transmitted to susceptible hosts. Moreover, urban areas may be susceptible to infectious disease spread due to human activities and host population densities [48, 49] which, when combined with our results, suggests the establishment of future pathogens anticipated under climate change may first occur, or impact of infection first be noted, in urban areas [2, 50].

## Supporting information

Sup figures and tables for Hanmer et al

## Acknowledgements

The authors would like to thank members of staff at APHA, BTO, ZSL and UKHSA who provided laboratory and technical support and Audra-Lynne Schlacter for undertaking IHC analysis on Blackbirds; members of the public who reported observations of garden bird mortality to the Garden Wildlife Health project (www.gardenwildlifehealth.org) and Animal and Plant Health Agency (APHA) and Scotland’s Rural College (SRUC) veterinary investigation centres for submitting birds for passive surveillance; staff and volunteers at Tiggywinkles Wildlife Hospital, Folly Wildlife Rescue and participating RSPCA wildlife centres for submitting Blackbirds that died or were euthanased in care for screening. In addition, we would like to thank Colin Wilson, Bill Haines, Tim Alexander, Martin Cade, BTO bird ringing groups and bird observatory staff and volunteers who have facilitated active sampling of passerines, all the participants in the BTO Garden BirdWatch who took part during the study period.

## Funding

This work was funded by a UKRI grant Vector-Borne RADAR (BB/X017990/1) and grant SV3045 from Department for Environment, Food & Rural Affairs (Defra) and the Devolved Administrations of Scotland and Wales. IoZ staff receive financial support from Research England. Financial support for the Garden Wildlife Health Project over the period 2023-2024 came from the Department for Environment, Food and Rural Affairs (Defra), the Welsh Government and the Animal and Plant Health Agency (APHA) Diseases of Wildlife Scheme (DoWS); and from the Garfield Weston Foundation and the Universities Federation for Animal Welfare. BTO Garden BirdWatch is funded by its participants through an annual subscription and additional donations, and would not be possible without their generosity and support.

## Author contributions

Conceptualisation: AAC, JMM, BL, RAR, AJF, Methodology: AAC, JMM, BL, RAR, AJF, Formal Analysis: HJH, RCB, BL, RAR, AJF, Investigation: All authors, Data Curation: HJH, RCB, AJJ, MS, AMI, BL, RAR, AJF, Writing-Original Draft: HJH, MS, RAR, BL, AJF, Writing Review & Editing: all authors, Visualisation: HJH, RAR, AJF, Supervision: JMM, BL, RAR, AJF, Project Administration: JMM, BL, RAR, AJF, Funding Acquisition: AAC, JMM, BL, RAR, AJF.

## Declaration of Interests

The authors declare no competing interests

## Supplementary information

All supplementary information is contained within journals supplementary files.

## Ethical Statement

All birds sampled through passive or enhanced passive surveillance were found dead or euthanised on welfare grounds. All active sampling of passerines described in the manuscript was licenced by the Home Office under project licence PP9908514: Identifying zoonotic flavivirus emergence pathways and seroprevalence in wild birds in the United Kingdom, and procedures were carried out by personal licence holders. Protocols were approved by the APHA Ethics Committee and the ZSL Ethics Committee (Animal Impacts) (IOZ202). All catching of birds was undertaken by trained ringers licenced through the British Trust for Ornithology following all relevant guidelines and regulations.

## Notes

### Competing Interest Statement

The authors have declared no competing interest.

