## Supplementary material for "Repeated and widespread introduction of a mosquito-borne virus, but wildlife impact remains localised in urban area": Sup figures and tables for Hanmer et al

**Supplementary files**

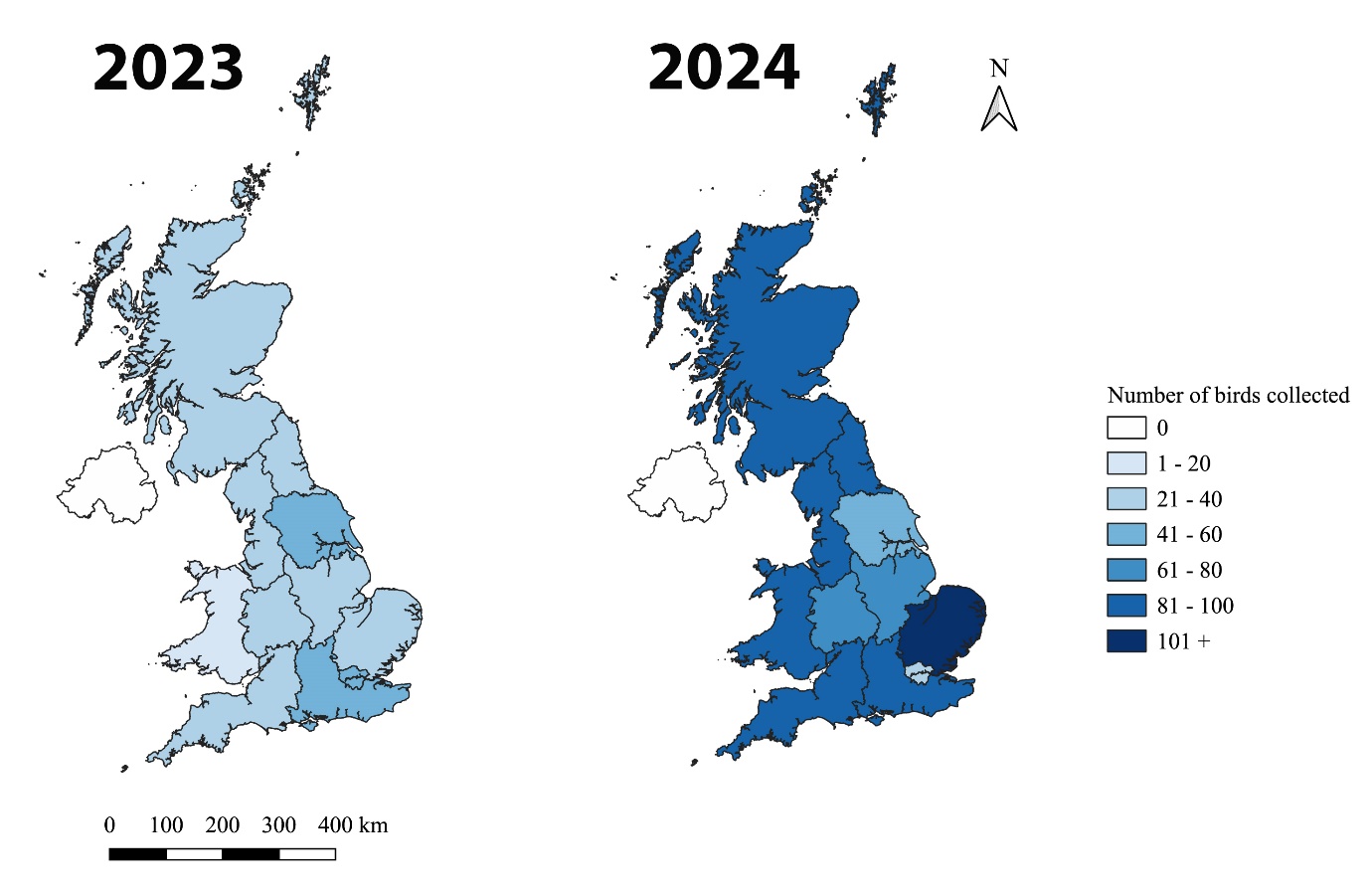

**Supplementary figure 1**. Passive surveillance in dead wild birds across the United Kingdom in 2023 (n = 559) (A) and 2024 (n =832) (B). Detailed breakdown of species and numbers below (supplementary table 1).

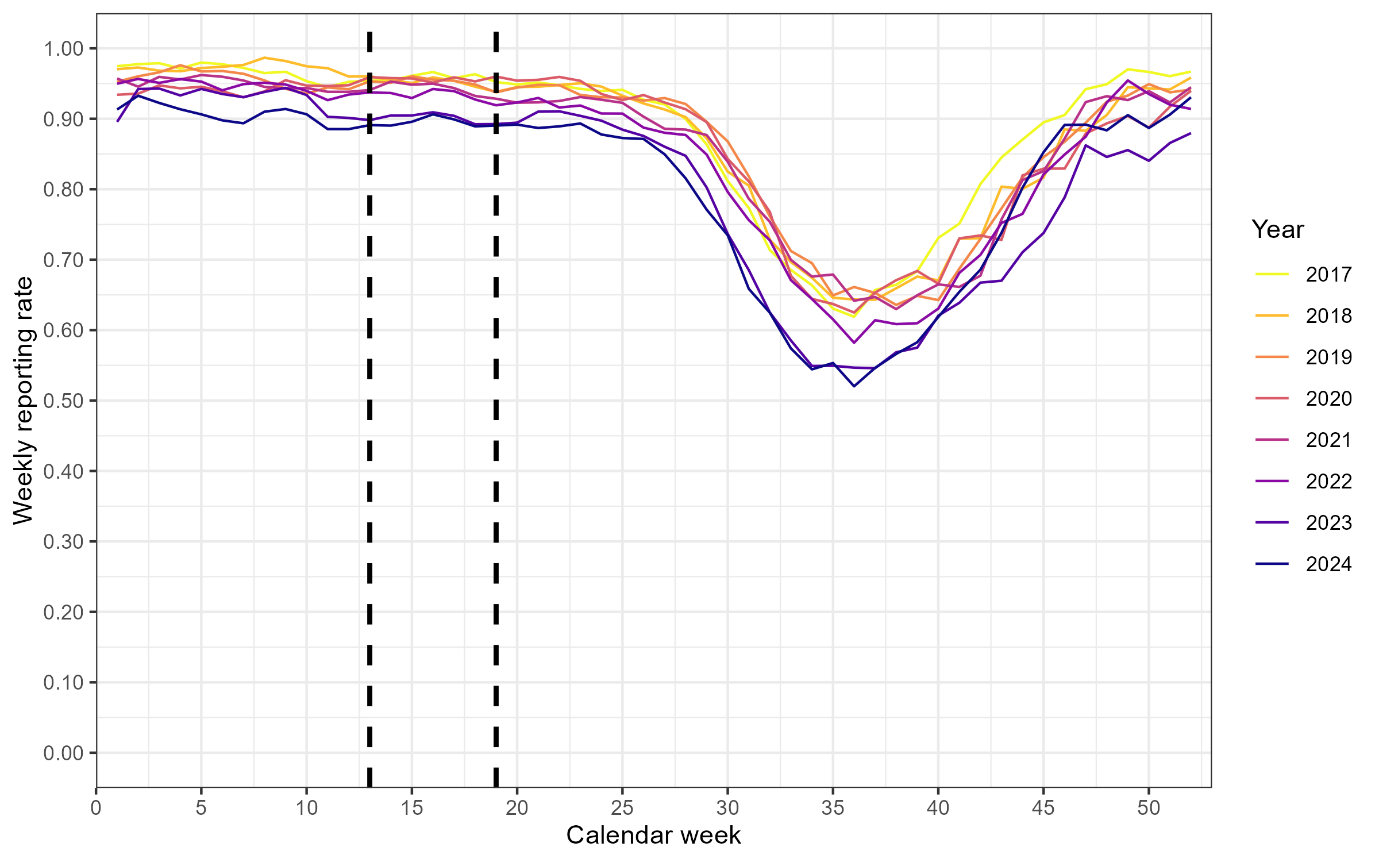

**Supplementary figure 2**. Weekly reporting rate for Blackbird in British Trust for Ornithology’s Garden BirdWatch across the UK for the period of 2017-2024 by year. Vertical dashed black lines indicate the first breeding attempt period defined as calendar weeks 13 to 19 inclusive (approximately 26^th^ March – 13^th^ May). All sites were required to be active for a minimum of two consecutive years with 67% coverage.

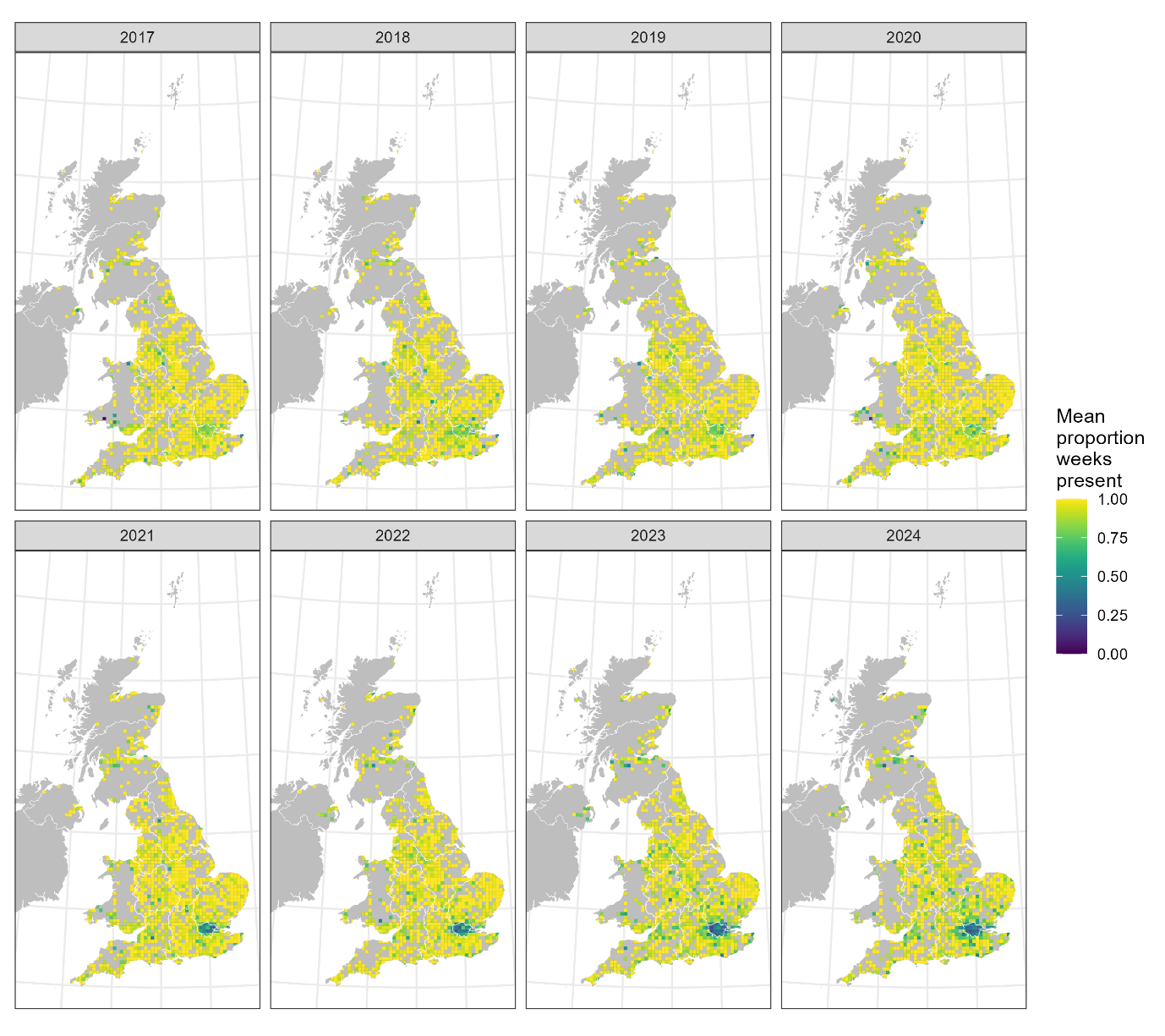

**Supplementary figure 3**. Maps showing change in the mean proportion of weeks when Blackbirds were recorded in British Trust for Ornithology’s Garden BirdWatch survey participant gardens per 10km grid square across the UK on an annual basis, over the period 2017-2024. Grey indicates less than two gardens fulfilling the criteria were active in that square in that year.

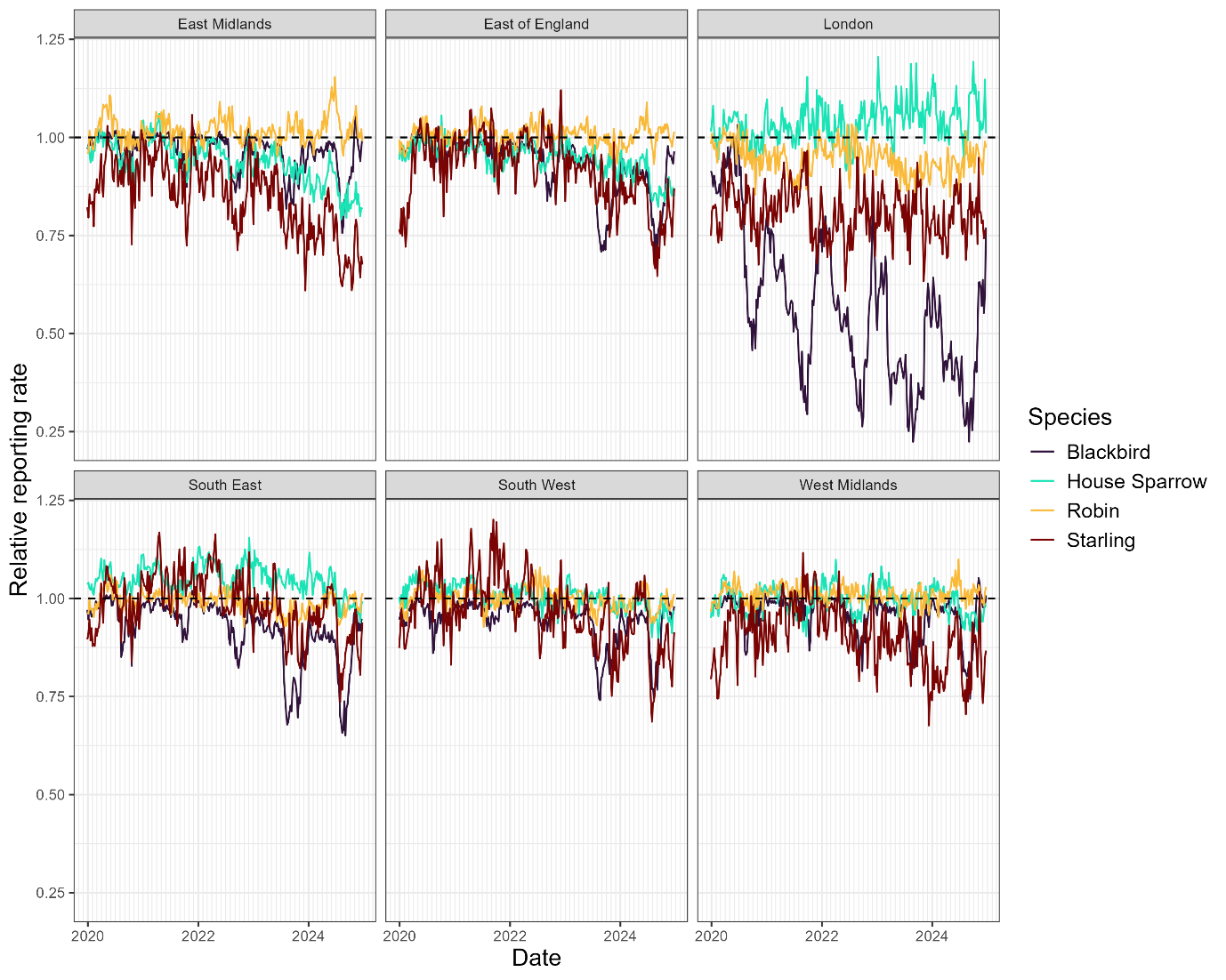

**Supplementary figure 4**. Regional relative weekly reporting rate for Eurasian Blackbird (*Turdus merula*), House Sparrow (*Passer domesticus*), Robin (*Erithacus rubecula*) and Starling (*Sturnus vulgaris*) over the period 2020-2024 inclusive. Data from British Trust for Ornithology’s Garden BirdWatch survey. A relative reporting rate of 1 indicates the presence of a species in a similar proportion of gardens in an assessed week compared to the 2010–2019 average for that week.

**Supplementary table 1**. Species composition of wild and captive birds (ɸ) tested for Usutu virus by RT-PCR, in 2023 (n=559) and 2024 (n=832).

| **Order** | **Family** | **Species** | **Number** | |
| --- | --- | --- | --- | --- |
|  |  |  | **2023** | **2024** |
| Accipitriformes | Accipitridae | Common Buzzard (*Buteo buteo*) | 40 | 37 |
|  |  | Eurasian Goshawk (*Astur gentilis*) | 14 | 3 |
|  |  | Golden Eagle (*Aquila chrysaetos*) | 2 | 3 |
|  |  | Hen Harrier (*Circus cyaneus*) | 4 | 10 |
|  |  | Honey Buzzard (*Pernis apivorus*) | 1 | - |
|  |  | Red Kite (*Milvus milvus*) | 10 | 15 |
|  |  | Sparrowhawk (*Accipiter nisus*) | 48 | 50 |
|  |  | White-Tailed Eagle (*Haliaeetus albicilla*) | - | 3 |
|  | Pandionidae | Osprey (*Pandion haliaetus*) | - | 5 |
| Anseriformes | Anatidae | Bewick’s Swan (*Cygnus columbianus*) | - | 1 |
|  |  | Canada Goose (*Branta canadensis*) | 40 | 59 |
|  |  | Egyptian Goose (*Alopochen aegyptiaca*) | - | 2 |
|  |  | Eider Duck (*Somateria mollissima*) | 1 | 1 |
|  |  | Greylag Goose (*Anser anser*) | 3 | 9 |
|  |  | Mallard (*Anas platyrhynchos*) | 17 | 26 |
|  |  | Mute Swan (*Cygnus olor*) | 47 | 95 |
|  |  | Pink-Footed Goose (*Anser brachyrhynchus*) | 2 | 1 |
|  |  | White-Faced Whistling Duck (*Dendrocygna viduata*) ɸ | 1 | - |
|  |  | Whooper Swan (*Cygnus cygnus*) | 1 | 3 |
| Bucerotiformes | Bucerotidae | Von der Decken’s Hornbill (*Tockus deckeni*) ɸ | 1 | - |
| Charadriiformes | Alcidae | Atlantic Puffin (*Fratercula arctica*) | 2 | 5 |
|  |  | Common Guillemot (*Uria aalge*) | 25 | 7 |
|  |  | Razorbill (*Alca torda*) | 15 | 1 |
|  | Charadriidae | Northern Lapwing (*Vanellus vanellus*) | - | 2 |
|  | Laridae | Arctic Tern (*Sterna paradisaea*) | - | 2 |
|  |  | Black-Headed Gull (*Chroicocephalus ridibundus*) | 5 | 2 |
|  |  | Black-Legged Kittiwake (*Rissa tridactyla*) | 1 | 6 |
|  |  | Common Gull (*Larus canus*) | 4 | 10 |
|  |  | Great Black-Backed Gull (*Larus marinus*) | 1 | 4 |
|  |  | Herring Gull (*Larus argentatus*) | 36 | 123 |
|  |  | Iceland Gull (*Larus glaucoides*) | 1 | - |
|  |  | Lesser Black-Backed Gull (*Larus fuscus*) | 7 | 20 |
|  |  | Mediterranean gull (*Ichthyaetus melanocephalus*) | 2 | - |
|  |  | Sandwich Tern (*Thalasseus sandvicensis*) | - | 1 |
|  |  | Yellow-Legged Gull (*Larus michahellis*) | - | 1 |
|  | Scolopacidae | Curlew (*Numenius arquata*) | - | 4 |
|  |  | Dunlin (*Calidris alpina*) | 3 | - |
|  |  | Eurasian Woodcock (*Scolopax rusticola*) | 1 | - |
| Ciconiiformes | Ciconiidae | Abdim’s Stork (*Ciconia abdimii*) ɸ | 2 | - |
|  |  | Woolly-Necked Stork (*Ciconia episcopus*) ɸ | - | 1 |
| Columbiformes | Columbidae | Black-Naped Fruit Dove (*Ptilinopus melanospilus*) ɸ | 1 | - |
|  |  | Collard Dove (*Streptopelia decaocto*) | 1 | 4 |
|  |  | Feral Pigeon (*Columba livia*) | 14 | 19 |
|  |  | Socorro Dove (*Zenaida graysoni*) ɸ | - | 1 |
|  |  | Wood Pigeon (*Columba palumbus*) | 11 | 13 |
| Falconiformes | Falconidae | Common Kestrel (*Falco tinnunculus*) | 7 | 13 |
|  |  | Peregrine Falcon (*Falco peregrinus*) | 4 | 3 |
| Galliformes | Phasianidae | Pheasent (*Phasianus colchicus*) | 7 | 9 |
|  |  | Red-Legged Partridge (*Alectoris rufa*) | - | 1 |
| Gaviiformes | Gaviidae | Great Northern Diver (*Gavia immer*) | - | 1 |
| Gruiformes | Rallidae | Eurasian Coot (*Fulica atra*) | 1 | 1 |
|  |  | Moorhen (*Gallinula chloropus*) | 2 | - |
| Musophagiformes | Musophagidae | Red-Crested Turaco (*Tauraco erythrolophus*) ɸ | 1 | - |
| Passeriformes | Corvidae | Carrion Crow (*Corvus corone*) | 11 | 25 |
|  |  | Common Raven (*Corvus corax*) | 4 | 1 |
|  |  | Magpie (*Pica pica*) | 2 | 7 |
|  |  | Jackdaw (*Coloeus monedula*) | 4 | 9 |
|  |  | Rook (*Corvus frugilegus*) | 2 | 14 |
|  | Emberizidae | Yellowhammer (*Emberiza citrinella*) | 3 | - |
|  | Estrildidae | Java Sparrow (*Padda oryzivora*) | 1 | - |
|  |  | Tricoloured Parrotfinch (*Erythrua tricolor*) ɸ | 1 | - |
|  | Fringillidae | Brambling (*Fringilla montifringilla*) | 1 | - |
|  |  | Bull Finch (*Pyrrhula pyrrhula*) | - | 2 |
|  |  | Chaffinch (*Fringilla coelebs*) | 8 | 5 |
|  |  | European Goldfinch (*Carduelis carduelis*) | 7 | 5 |
|  |  | Greenfinch (*Carduelis chloris*) | 10 | 14 |
|  |  | Siskin (*Carduelis spinus*) | 1 | 3 |
|  | Hirundinidae | House Martin (*Delichon urbicum*) | - | 1 |
|  | Leiothrichidae | Emei Shan Liocichla *(Liocichla omeiensis)* ɸ | 1 | - |
|  |  | Red-Tailed Laughingthrush (*Trochalopteron milnei*) ɸ | - | 1 |
|  |  | Silver-Eared Mesia (*Leiothrix argentauris*) ɸ | - | 1 |
|  |  | Sumatran Laughingthrush (*Garrulax bicolor*) ɸ | - | 1 |
|  | Muscicapidae | Nightingale (*Luscinia megarhynchos*) | 1 | - |
|  |  | European Robin (*Erithacus rubecula*) | 1 | 1 |
|  |  | Spotted flycatcher (*Muscicapa striata*) | 1 | - |
|  |  | White-Crowned Robin-Chat (*Cossypha albicapillus*) ɸ | 1 | - |
|  | Paridae | Blue Tit (*Cyanistes caeruleus*) | - | 3 |
|  |  | Coal Tit (*Periparus ater*) | 1 | - |
|  |  | Great Tit (*Parus major*) | 1 | - |
|  | Passeridae | House Sparrow (*Passer domesticus*) | 11 | 5 |
|  | Prunellidae | Dunnock (*Prunella modularis*) | 1 | 1 |
|  | Regulidae | Goldcrest (*Regulus regulus*) | - | 2 |
|  | Sturnidae | Common Starling (*Sturnus vulgaris*) | 9 | 7 |
|  |  | Grosbeak Starling (*Scissirostrum dubium*) ɸ | - | 2 |
|  | Sylviidae | Blackcap (*Sylvia atricapilla*) | 1 | - |
|  | Turdidae | Blackbird (*Turdus merula*) | 16 | 68 |
|  |  | Chestnut-Backed Thrush (*Geokichla dohertyi*) ɸ | - | 4 |
|  |  | Song Thrush (*Turdus philomelos*) | 10 | 2 |
|  | Troglodytidae | Wren (*Troglodytes troglodytes*) | - | 3 |
| Pelecaniformes | Ardeidae | Grey Heron (*Ardea cinerea*) | - | 5 |
|  | Threskiornithidae | Eurasian Spoonbill (*Platalea leucorodia*) | 1 | - |
|  |  | Waldrapp Ibis (*Geronticus eremita*) ɸ | 1 | - |
| Piciformes | Picidae | Great Spotted Woodpecker (*Dendrocopos major*) | 3 | 2 |
|  |  | Green Woodpecker (*Picus viridis*) | 1 | - |
| Podicipediformes | Podicipedidae | Great Crested Grebe (*Podiceps cristatus*) | - | 1 |
| Procellariiformes | Procellariidae | Manx Shearwater (*Puffinus puffinus*) | - | 1 |
| Sphenisciformes | Spheniscidae | Black-Footed Penguin (*Spheniscus demersus*) ɸ | 3 | - |
|  |  | Humboldt Penguin (*Spheniscus humboldti*) | 8 | 1 |
| Strigiformes | Strigidae | Great Grey Owl (*Strix nebulosa*) ɸ | 2 | - |
|  |  | Little Owl (*Athene noctua*) | 1 | - |
|  |  | Long-Eared Owl (*Asio otus*) | 1 | - |
|  |  | Tawny Owl (*Strix aluco*) | 25 | 25 |
|  | Tytonidae | Barn Owl (*Tyto alba*) | 12 | 17 |
| Suliformes | Phalacrocoracidae | European Shag (*Gulosus aristotelis*) | - | 2 |
|  |  | Great Cormorant (*Phalacrocorax carbo*) | 3 | 7 |
|  | Sulidae | Northern Gannet (*Morus bassanus*) | 2 | 6 |
| Trogoniformes | Trogonidae | Collared Trogon (*Trogon collaris*) ɸ | - | 1 |

**Supplementary table 2**. Usutu virus whole genome sequences obtained from Genbank and used in phylogenetic analysis. All sequences used from GenBank, aside from ones of UK origin, are detections of USUV made in the Netherlands as these represent a likely route of incursion to the UK.

| **GenBank accession number** | **Usutu lineage** | **Sequence Length (base pairs)** |
| --- | --- | --- |
| MN122238 | Africa 3.1 | 10932 |
| MN122236 | Africa 3.2 | 10932 |
| MN122201 | Africa 3.2 | 10932 |
| MN122163 | Africa 3.2 | 10932 |
| MN122145 | Africa 3.2 | 10932 |
| MN122249 | Africa 3.2 | 10932 |
| MN122238 | Africa 3.2 | 10932 |
| MN122148 | Africa 3.2 | 10932 |
| MN122151 | Africa 3.2 | 10932 |
| MN122156 | Africa 3.2 | 10932 |
| MN122162 | Africa 3.2 | 10932 |
| MW001216 (UK) | Africa 3.2 | 10922 |
| OM202464 (UK) | Africa 3.2 | 10922 |
| MN122167 | Africa 3.3 | 10932 |
| MN122168 | Africa 3.3 | 10932 |
| MN122205 | Africa 3.3 | 10932 |
| MN122234 | Africa 3.3 | 10932 |
| MN122253 | Africa 3.3 | 10932 |
| MN122237 | Africa 3.3 | 10932 |
| MN122239 | Africa 3.3 | 10932 |

**Supplementary table 3**. Wild and zoological collection birds tested through enhanced passive and active surveillance, that were RT-PCR positive for Usutu virus (n = 23) or were seropositive for anti-*Orthoflavivirus* antibodies using an ELISA (n =8) and/or USUV specific antibodies via plaque reduction neutralisation test (PRNT) (n=4) in chronological order over the period July 2023-December 2024, inclusive. N/A=Not applicable.

| **Species & unique identifier** | **Life stage** | **Sample** | **Surveillance type** | **Date of collection (active sampling) or death (passive sampling)** | **Location** | **PCR result and lineage** | **ELISA** | **PRNT** |
| --- | --- | --- | --- | --- | --- | --- | --- | --- |
| Blackbird (*Turdus merula*)*  XT0497-23 | Juvenile | Pooled brain & kidney | Passive | 17/07/2023 | Greater London | Positive  Africa 3.2 (tissue *Ct* 26.9) | N/A | N/A |
| Blackbird (*Turdus merula*)  XT0504-23 | Juvenile | Pooled brain & kidney | Passive | 24/07/2023 | Cambridgeshire | Positive  Africa 3.2 (tissue *Ct* 25.1) | N/A | N/A |
| Feral pigeon (*Columba livia*)*  XT0624-23 | Adult male | Pooled brain & kidney | Passive | 11/09/2023 | Greater London | Positive  Africa 3.2 (tissue *Ct* 30.9) | N/A | N/A |
| Blackbird (*Turdus merula*) *  XT0630-23 | Juvenile | Pooled brain & kidney  Primary feather | Passive | 12/09/2023 | Greater London | Positive  Africa 3.2  (tissue *Ct* 23.72  feather no *Ct*) | N/A | N/A |
| Blackbird (*Turdus merula*)  XT0078-24 | Adult male | Pooled brain & kidney  Primary feather | Passive – Wildlife centre sentinel | 13/09/2023 | Buckinghamshire | Positive  Africa 3.2  (tissue *Ct* 21  feather *no Ct*) | N/A | N/A |
| Blackbird (*Turdus merula*)  XT0122-24 | Adult male | Pooled brain & kidney  Primary feather | Passive – Wildlife centre sentinel | 14/09/2023 | Bedfordshire | Positive  Africa 3.2  (tissue C*t* 23  feather no *Ct*) | N/A | N/A |
| Great grey owl (*Strix nebulosa*)* ɸ  ZB0631-23 | Unknown | Pooled brain & kidney | Passive – Zoo collection animal | 14/09/2023 | Greater London | Positive  Africa 3.2  (tissue *Ct* 22.42) | N/A | N/A |
| Great grey owl (*Strix nebulosa*)* ɸ  ZB0634-23 | Unknown | Pooled brain & kidney | Passive – Zoo collection animal | 16/09/2023 | Greater London | Positive  Africa 3.2  (tissue *Ct* 24.51) | N/A | N/A |
| Blackbird (*Turdus merula*)  XT0088-24 | Juvenile male | Pooled brain & kidney | Passive – Wildlife centre sentinel | 28/09/2023 | Oxfordshire | Positive  Africa 3.2  (tissue *Ct* 34) | N/A | N/A |
| Blackbird (*Turdus merula*)  XT0021-24 | Adult male | Pooled brain & kidney  Primary feather | Passive – Wildlife centre sentinel | 01/10/2023 | East Sussex | Positive  Africa 3.1  (tissue *Ct* 23  feather no *Ct*) | N/A | N/A |
| Blackbird (*Turdus merula*)  XT0691-23 | Adult male | Pooled brain & kidney | Passive | 09/10/2023 | Cambridgeshire | Positive  Africa 3.2  (tissue *Ct* 26.1) | N/A | N/A |
| Whitethroat (*Curruca communis*) | Juvenile | Feather | Active | 22/07/2024 | Hampshire | Positive  Africa 3.2  (tissue *Ct* 34.0) | Negative | N/A |
| Chiffchaff (*Phylloscopus collybita*) | Juvenile | Blood | Active | 22/07/2024 | Hampshire | Negative | Positive | Insufficient serum |
| Whitethroat (*Curruca communis*) | Juvenile | Blood | Active | 22/07/2024 | Hampshire | Negative | Positive | Insufficient serum |
| Blackbird (*Turdus merula*) | Juvenile | Blood | Active | 06/08/2024 | Kent | Negative | Positive | Insufficient serum |
| Willow Warbler (*Phylloscopus trochilus*) | Juvenile | Blood | Active | 06/08/2024 | Kent | Negative | Positive | Insufficient serum |
| Blackbird (*Turdus merula*)  XT0411-25 | Adult male | Pooled brain & kidney  Primary feather | Passive – Wildlife centre sentinel | 23/08/2024 | Hertfordshire | Positive  Africa 3.2  (tissue *Ct* 30.6  Feather *Ct* 34.1) | N/A | N/A |
| Grosbeak Starling (*Scissirostrum dubium*)* ɸ | Unknown | Pooled brain & kidney | Passive – Zoo collection animal | 02/09/2024 | Greater London | Positive  Africa 3.2  (tissue *Ct* 25.52) | N/A | N/A |
| Grosbeak Starling (*Scissirostrum dubium*)* ɸ | Unknown | Pooled brain & kidney | Passive – Zoo collection animal | 02/09/2024 | Greater London | Positive  Africa 3.2  (tissue *Ct* 29.55) | N/A | N/A |
| Blackbird (*Turdus merula*)  XT0321-25 | Adult female | Pooled brain & kidney  Primary feather | Passive – Wildlife centre sentinel | 09/09/2024 | Buckinghamshire | Positive  Africa 3.2  (tissue *Ct* 28.7  Feather *Ct* 33.2) | N/A | N/A |
| Chiffchaff (*Phylloscopus collybita*) | Juvenile | Contour feather | Active | 21/09/2024 | Cambridgeshire | Positive  Africa 3.2 (feather Ct 34.9) | Negative | N/A |
| Blackcap (*Sylvia atricapilla*) | Juvenile | Feather | Active | 10/10/2024 | Dorset | Positive  Africa 3.2  (feather *Ct* 20.84) | Negative | N/A |
| Blackcap (*Sylvia atricapilla*) | Juvenile | Cloacal swab | Active | 10/10/2024 | Dorset | Positive  Africa 3.2  (swab *Ct* 33.59) | Negative | N/A |
| Chiffchaff (*Phylloscopus collybita*) | Juvenile | Feather | Active | 10/10/2024 | Dorset | Positive  Africa 3.2  (feather *Ct* 35.65) | Negative | N/A |
| Chiffchaff (*Phylloscopus collybita*) | Juvenile | Feather | Active | 10/10/2024 | Dorset | Positive  Africa 3.2  (feather *Ct* 30.76) | Negative | N/A |
| Blackcap (*Sylvia atricapilla*) | Juvenile | Feather | Active | 15/10/2024 | Kent | Positive  Africa 3.2  (feather *Ct* 32.42) | Negative | N/A |
| Chiffchaff (*Phylloscopus collybita*) | Juvenile | Oral swab | Active | 15/10/2024 | Kent | Positive  Africa 3.2  (swab *Ct* 33.85) | Negative | N/A |
| Blackbird (*Turdus merula*) | Adult | Blood | Active | 11/12/2024 | Berkshire | Negative | Positive | USUV neutralising antibodies detected |
| Blackbird (*Turdus merula*) | Adult | Blood | Active | 11/12/2024 | Berkshire | Negative | Positive | USUV neutralising antibodies detected |
| Blackbird (*Turdus merula*) | Adult | Blood | Active | 13/12/2024 | Hampshire | Negative | Positive | USUV neutralising antibodies detected |
| Blackbird (*Turdus merula*) | Adult | Blood | Active | 13/12/2024 | Hampshire | Negative | Positive | USUV neutralising antibodies detected |

* Birds recovered from UK USUV index site (ZSL London Zoo, Greater London)

ɸ Collection birds from ZSL London Zoo, Greater London

**Supplementary table 4**. Wild bird post-mortem examination and ancillary diagnostic findings of Usutu virus (USUV) RT-PCR-positive wild birds.

Bursa of Fabricius (BoF); BW=Body weight; GIT=Gastrointestinal tract; NA=Not available; NAD=No abnormalities detected; ND=Not done; NSF=No significant findings; SI=Small intestine.

Key macroscopic findings include summary description of liver and spleen, including organ weight and dimensions, and skin cranial to the vent, given published observations of hyperkeratosis, ulceration and sero-cellular crusts in this area in association with USUV infection (Giglia *et al.* 2021. *Viruses*. **13**:1481. doi: [10.3390/v13081481](https://doi.org/10.3390/v13081481)).

Immunohistochemical labelling against flavivirus envelope (E) antigen classification: ND not done; (-) absent; (+) rare; + mild; ++ moderate; +++ abundant.

| Species & unique Identifier | History | Signalment & body condition & BW (g) & state of carcase preservation | Key macroscopic findings | Microbiology | Parasitology | Cytology/Histopathology | Immunohistochemistry |
| --- | --- | --- | --- | --- | --- | --- | --- |
| Blackbird  XT0497-23 | Found dead. | Juvenile, undetermined sex.  Normal body condition. 87.5 g  Fresh.  Mild autolysis. | Left lung diffusely red.  GIT with a moderate volume of black, liquid to paste-like content.  **Liver:** NSF (weight NA).  **Spleen:** splenomegaly (40 x 10 x 7 mm; weight NA)  **Skin cranial to the vent:** NA. | **SI contents:** NSF.  **Spleen:** NSF. | **SI contents:** Numerous adult acanthocephalans & scant *Strongyle*-type ova. | Brain cytology: several protozoal schizonts (suspected *Plasmodium* sp.) within the endothelium of capillaries. | ND |
| Blackbird  XT0504-23 | Weak, orphaned.  Died. | Juvenile, undetermined sex.  Normal body condition. 61.7 g  Fresh. Moderate autolysis. | Lungs dark red.  Gizzard with scant contents (no recent feeding).  Kidneys red, friable, with uneven surface (suspected due to autolysis).  **Liver**: dark brown to red (3.7 g).  **Spleen:** dark red (17 x 4.5 x 4 mm; weight NA).  **Skin cranial to the vent:** NSF. | **Liver:** NSF.  **SI contents:** NSF. | **SI contents:** Numerous adult cestodes. | ND | ND |
| Feral pigeon  XT0624-23 | Unable to fly.  Euthanasia.  . | Adult male.  Thin.  Weight NA  Fresh.  Minimal autolysis. | Proliferative skin lesions on face (suspected avian pox). | NA. | **SI contents**: Scant protozoal oocysts. | ND | ND |
| Blackbird  XT0630-23 | Lethargic, unable to fly, gasping.  Died. | Juvenile, undetermined sex.  Emaciated. 76.8 g  Fresh.  Moderate autolysis. | Skin adherent to subcutis (suspected dehydration).  Lungs very dark red, wet and frothy.  Gizzard empty (no recent feeding).  **Liver**: very dark red, sharp lobe margins, extending beyond keel, equivocal hepatomegaly (6.5 g).  **Spleen**: very dark red, splenomegaly (39.3 x 6.7 x 6.7 mm; 1.1 g)  **Skin cranial to the vent**: small area with crusts present (dimensions NA). | **Liver**: Confluent nearly pure isolate *Escherichia coli* 1. with *Staphylococcus* sp. (aurex -neg).  **SI contents**: NSF. | **SI contents**: Adult Ascaridae. | **Heart:** Minimal multifocal myocardial necrosis.  **Lungs:** Generalised congestion.  **Liver, spleen:** Moderate multifocal necrosis.  **Skin cranial to the vent:** Moderate multifocal subacute lymphocytic and heterophilic dermatitis.  BoF, brain, skeletal muscle, trachea: NSF.  GIT and kidney: Moderate autolysis precluding meaningful interpretation. | (-) SI  (+) Cerebellum, heart, kidney, lung, skeletal muscle, trachea  + BoF, gizzard, liver, proventriculus, skin cranial to the vent  +++ Spleen |
| Blackbird  XT0078-24 | Unable to stand or fly.  Died. | Adult male.  Emaciated. 64.6 g  Frozen.  Moderate autolysis. | Scant contents throughout GIT (no recent feeding).  **Liver**: dark red, slightly friable, normal to small size (2.3 g).  **Spleen**: dark red (29.5 x 5.6 x 5.6 mm; 0.3 g).  **Skin cranial to the vent**: area of scale present (8 x 6 mm). | **Liver**: NSF.  **SI contents**: NSF. | **SI contents**: Moderate *Raillietina* sp. ova. | **Cerebral cortex**: Mild oligofocal acute neuronal necrosis.  **Heart**: Minimal multifocal myocardial necrosis.  **Skin cranial to the vent**: Mild multifocal lymphocytic perivascular dermatitis, with focal dermal oedema and pustular formation.  Gizzard, kidney, liver, lung, proventriculus, SI, skeletal muscle, spleen, trachea: NSF. | (+) Lung  + Cerebellum, gizzard, proventriculus, SI, spleen  ++ Cerebrum, heart, kidney, liver, skin cranial to the vent, trachea |
| Blackbird  XT0122-24 | Unable to fly.  Died. | Adult male.  Thin. 91.3 g  Frozen.  Moderate autolysis. | Skin adherent to subcutis (suspected dehydration).  Gizzard with scant contents (no recent feeding).  **Liver**: dark red-brown, sharp lobe margins (4.4 g). **Spleen**: plum-coloured, equivocal splenomegaly  (50.1 x 5.1 x 5.0 mm; 0.6 g).  **Skin cranial to the vent**: : area of scale present (approximately 5 x 5 mm). | **Liver**: NSF.  **SI contents**: NSF. | **SI contents**: Moderate adult Ascaridae, numerous *Raillietina* sp. ova. | **Skin cranial to the vent**: Mild focal acute pustular dermatitis, with marked multifocal to coalescing monocytic and granulocytic perivascular infiltrates.  BoF, brain, gizzard, heart, kidney, liver, lung, oesophagus, SI, skeletal muscle, spleen, trachea: NSF. | (-) SI  (+) Skeletal muscle  + Kidney, trachea  ++ Gizzard, liver, lung, spleen  +++ Heart, skin cranial to the vent |
| Blackbird  XT0088-24 | Unable to fly.  Died. | Juvenile male.  Thin. 90.1 g  Frozen.  Mild autolysis. | Trauma (fracture of left femur).  Bilateral adrenomegaly.  Gizzard empty (no recent feeding). Intestinal serosa has a mixed colouration, with some very dark red patches (suspected haemorrhage).  **Liver**: dark brown-black, friable, sharp lobe margins but extending beyond the limits of the keel, equivocal hepatomegaly (6.5 g).  **Spleen**: dark red, homogenous, plump appearance, splenomegaly (39 x 9 x 9 mm; 1.0 g).  **Skin cranial to the vent**: : area of scale present (7 x 6 mm). | **Liver**: NSF.  **SI contents**: NSF. | **SI contents**: Scant to moderate protozoal oocysts & scant to moderate adult nematodes. | Brain, heart, kidney, liver, lung, oesophagus, trachea, SI, skeletal muscle, skin cranial to the vent, spleen: moderate to marked autolysis; NSF. | (-) Cerebrum, gizzard, heart, kidney, liver, lung, oesophagus, SI, skeletal muscle, skin cranial to the vent, spleen, trachea |
| Blackbird  XT0021-24 | Unable to fly.  Euthanasia. | Adult male.  Thin. 74.0 g  Frozen.  Mild autolysis. | Two *Ixodes frontalis* ticks on submandibular region, not engorged and no visible reaction.  Left peri-ocular swelling, dark red conjunctiva.  Right lung dark red, wet and frothy.  Gizzard empty (no recent feeding).  **Liver**: sharp lobe margins, but liver extending beyond limit of keel (equivocal hepatomegaly, however possible artifact due to intra-hepatic euthanasia) (weight NA).  **Spleen**: dark red, equivocal splenomegaly  (42 x 5 x 5 mm; weight NA).  **Skin cranial to the vent**: thickened area (7 x 8 mm). | **Liver**: NSF.  **SI contents**: NSF. | **SI contents**: Numerous *Raillietina* sp. ova & moderate adult nematodes & numerous larval helminths. | **Eye**: Mild multifocal granulocytic and lymphocytic periocular dermatitis (most likely related to peri-ocular trauma noted at postmortem).  **Trachea, gizzard and SI**: Mild to moderate multifocal lymphocytic perivascular infiltrates.  **Liver**: Minimal random multifocal hepatocellular degeneration.  **Skin cranial to the vent**: Moderate focal acute ulcerative dermatitis with mild multifocal dermal monocytic infiltrates.  Brain, kidney, heart, kidney, large intestine, lung, proventriculus, SI, skeletal muscle, spleen: NSF. | (-) Cerebrum  (+) Lung, skeletal muscle  + Gizzard, oesophagus, trachea  ++ Eye, heart, kidney, skin cranial to the vent  +++ Liver, SI, spleen |
| Blackbird  XT0691-23 | Found dead. | Adult male.  Emaciated.  83.0 g  Fresh.  Moderate autolysis. | Scant contents throughout GIT (no recent feeding).  **Liver**: dark red-brown, sharp lobe margins, equivocal hepatomegaly (5.0 g).  **Spleen**: dark red-black, equivocal splenomegaly (43.0 x 7.5 x 5.5 mm; weight NA).  **Skin cranial to the vent**: NAD. | **Liver**: NSF.  **SI contents**: NSF. | **SI contents**: Negative. | **Liver:** Mild multifocal necrosis.  **Spleen:** Mild multifocal necrosis.  **Skin from head:** Mild multifocal monocytic dermal vasculitis with prominent mononuclear perivascular and perineural infiltrates.  Brain, gizzard, heart, kidney, large intestine, lung, oesophagus, SI, skeletal muscle, trachea: NSF.  All: mild to moderate autolysis. | (+) Trachea  + Brain, gizzard, kidney, lung, skeletal muscle  ++ SI  +++ Heart, liver, skin from head, spleen |
| Blackbird  XT0411-25 | Ataxia.  Died. | Adult male.  Emaciated.  58.4 g  Frozen.  Moderate autolysis. | Vent heavily soiled with urates.  Gizzard empty (no recent feeding).  **Liver**: dark plum-brown, sharp lobe margins, normal to small size (2.1 g).  **Spleen**: dark red (20.6 x 3.6 x 2.6 mm; 0.2 g).  **Skin cranial to the vent**: thickened, with scale and crusts in equal proportion (12 x 8 mm affected). | **Liver**: NSF.  **SI contents**: NSF. | **SI contents**: Scant adult acanthocephalans. | **Heart:** Moderate multifocal monocytic myocarditis.  **Liver:** Mild multifocal monocytic periportal hepatitis.  **Gizzard, small intestine, large intestine:** Mild to moderate mononuclear perivascular infiltrates observed in the serosa.  **Skin cranial to the vent:** Moderate multifocal monocytic dermatitis with marked multifocal monocytic perivascular infiltrates (dermis and subcutis).  Brain, kidney, lung, oesophagus, skeletal muscle, trachea: NSF.  All: mild to moderate autolysis, freeze-thaw artefact. | + Kidney, liver, lung, SI, spleen  ++ Gizzard, heart  +++ Cerebrum, skin cranial to the vent |
| Blackbird  XT0321-25 | Wounds to body, presence of ticks.  Died. | Adult female.  Thin.  66.4 g  Frozen.  Moderate autolysis. | Polytrauma (fracture of left ulna and keel).  Scant contents throughout GIT (no recent feeding).  **Liver**: dark plum, sharp lobe margins (3.5 g).  **Spleen**: dark plum (24 x 5 x 5 mm; 0.3 g).  **Skin cranial to the vent**: crusts present, with a smaller proportion of scale (9 x 8 mm affected). | **Liver**: NSF.  **SI contents**: NSF. | **SI contents**: Adult cestodes, protozoal oocysts. | **Lungs:** Minimal multifocal monocytic interstitial pneumonia.  **Heart:** Minimal multifocal lymphohistiocytic epicarditis and myocarditis with multifocal moderate lymphohistiocytic perivascular and perineural infiltrates.  **Oesophagus, gizzard, small intestine, large intestine:** Mild to moderate mononuclear perivascular infiltrates.  **Skeletal muscle:** Moderate multifocal necrotising myositis.  **Skin cranial to the vent:** Moderate multifocal monocytic dermatitis with multifocal marked monocytic perivascular infiltrates.  Brain, kidney, trachea: NSF.  Liver and spleen: Moderate autolysis and freeze-thaw artefact precluding meaningful interpretation. | (-) Liver, trachea  + Kidney, lung, oesophagus, skeletal muscle  ++ Cerebrum, gizzard, heart, SI, skin cranial to the vent, spleen |
